# Evolutionary origin of synovial joints

**DOI:** 10.1101/2024.04.02.587820

**Authors:** Neelima Sharma, Yara Haridy, Neil Shubin

**Affiliations:** Department of Organismal Biology and Anatomy, The University of Chicago, Chicago, IL, USA

## Abstract

Synovial joints, characterized by reciprocally congruent and lubricated articular surfaces separated by a cavity, are hypothesized to have evolved from continuous cartilaginous joints for increased mobility and improved load bearing. To test the evolutionary origins of synovial joints, we examine the morphology, genetic, and molecular mechanisms required for the development and function of the joints in elasmobranchs and cyclostomes. We find the presence of cavitated and articulated joints in elasmobranchs, such as the little skate (*Leucoraja erinacea*) and bamboo shark (*Chiloscyllium plagiosum*), and the expression of lubrication-related proteoglycans such as aggrecan and glycoproteins such as hyaluronic acid receptor (CD44) at the articular surfaces in little skates. Sea lampreys (*Petromyozon marinus*), a representative of cyclostomes, are devoid of articular cavities but express proteoglycan-linking proteins throughout their cartilaginous skeleton, suggesting that the expression of proteoglycans is primitively not limited to the articular cartilage. Analysis of the development of joints in the little skate reveals the expression of growth differentiation factor-5 (*Gdf5*) and *β*-catenin at the joint interzone before the process of cavitation, indicating the involvement of BMP and Wnt-signaling pathway, and reliance on muscle contraction for the process of joint cavitation, similar to tetrapods. In conclusion, our results show that synovial joints are present in elasmobranchs but not cyclostomes, and therefore, synovial joints originated in the common ancestor of extant gnathostomes. A review of fossils from the extinct clades along the gnathostome stem further shows that synovial joints likely arose in the common ancestor of gnathostomes. Our results have implications for understanding how the evolution of synovial joints around 400 mya in our vertebrate ancestors unlocked motor behaviors such as feeding and locomotion.

**Author summary:** We owe our mobility and agility to synovial joints, characterized by a lubricated joint cavity between the bony elements. Due to the cavity, synovial joints function by bones sliding relative to each other, allowing an extensive range of motion and heightened stability compared to fused or cartilaginous joints that function by bending. Using histological and protein expression analysis, we show that reciprocally articulated, cavitated, and lubricated joints are present in elasmobranchs such as skates and sharks but not in cyclostomes such as the sea lamprey. Furthermore, the development of the little skate joints relies on genetic regulatory mechanisms such as BMP and Wnt-signalling, similar to tetrapods. Thus, our results show that synovial joints are present in elasmobranchs but not in cyclostomes. In conclusion, synovial joints originated in the common ancestor of jawed vertebrates. Furthermore, a review of fossil taxa along the gnathostome stem shows that cavitated joints that function by relative sliding of articulating surfaces originated at the common ancestor of all gnathostomes. Our results have consequences for understanding how the evolution of cavitated and lubricated joints in ancient vertebrates impacted behaviors like feeding and locomotion 400 million years ago.

## Introduction

Synovial joints are an innovation that allowed vertebrates to have improved load-bearing capacity, a greater range of motion, better control of rotational degrees of freedom, and stability [1]. Comparative and molecular analyses shows the presence of synovial-like joint morphology in osteichthyans [2] because zebrafish, gar, and lungfish exhibit lubricated and cavitated joints and studies place the origin of synovial joints at the common ancestor of bony fishes. However, we know relatively little about the characteristics and development of joints in the clades that phylogenetically precede osteichthyans, such as chondrichthyans and cyclostomes, and extinct taxa phylogenetically between them. Therefore, when synovial joints originated in vertebrates remains uncertain.

Synovial joints consist of a characteristic lubricated joint cavity that separates the bones, and each bony element is covered by a thin layer of hyaline articular cartilage [3]. Tetrapods [4] and bony fishes [2] exhibit articular cartilage separated into superficial, transitional, middle, and calcified zones, which separate the radial uncalcified articular cartilage from the subchondral bone [5]. Shared expression of chondrocytes and extracellular matrix (ECM) proteins such as collagen-I and collagen-II along with the expression of proteoglycans such as aggrecan and lubricin, glycosaminoglycans (GAGs) such as hyaluronic acid, and glycoproteins such as CD44 at the articular regions suggest synovial joint morphology [4, 6]. Aggrecan, lubricin, and other lubricating proteins are secreted in the joint cavity to provide a near frictionless joint motion and prevent the chondrocytes and articular surfaces from damage [2].

The development of synovial joints relies on the emergence of an interzone in the uninterrupted mesenchyme that cavitates to give rise to articular surfaces and joint cavity. Molecular mechanisms underlying synovial joint formation involve the activity of BMP and the Wnt signaling pathways. Growth differentiation factor 5 (*Gdf5*), related to the BMP signaling pathway, is continuously expressed by an influx of newly differentiated chondrocytes at the interzone in the uninterrupted cartilage [7]. Gdf5+ cells correlate with the subsequent expression of Wnt genes, with *β*-catenin as the central player. Joint cells expressing *β*-catenin are the joint cell progenitors that give rise to articular cartilage, capsule, and ligaments, and loss of *β*-catenin causes joint fusion [8]. Canonical Wnt-signaling maintains the subsequent fate of the joint interzone cells but it does not play a role in the induction of the interzone [8–10]. Apart from molecular mechanisms of morphogenesis, muscle contraction is often necessary for bone elongation and joint cavity formation [11]. The process of cavitation relies on skeletal extension and flexion that causes the formation and expansion of microcavities in the interzone [12]. In the absence of muscles or muscle contraction, Wnt-signaling is weaker [13], *Gdf5* expression is lost [11], skeletal segments are smaller compared to controls [14, 15], and the joints are fused [11, 16].

The literature on the joints of chondrichthyans and cyclostomes is scarce, but studies have addressed cartilage and jaw evolution in vertebrates by comparing developmental and molecular mechanisms in jawless lampreys with jawed chondrichthyans. Both lamprey and chondrichthyan cartilage comprises of collagen-II [17–19], generated by similar underlying genetic regulatory network [20]. During development, sea lampreys express *Gdf* in the pharyngeal arches and more generally in the developing skeleton, but not in the intermediate region of the first arch [21, 22]. *Gdf5* recruitment in the intermediate first arch is hypothesized to have led to the evolution of jaws in gnathostomes [21, 23]. Indeed, little skate expresses gdf5 in the intermediate region of all the arches [24]; however, its expression at the interzone in the rest of the skeleton at the later stages of embryogenesis is unclear. Notably, Davies, 1948 [25] described synovial joints in skates based on dissection studies that showed a joint cavity, ligaments, and synovial membrane. However, whether synovial joints are a common feature in elasmobranchs and whether they are developmentally and molecularly similar to osteichthyans remains unknown.

In this study, we test when synovial joints arose in evolution by analyzing the joints of cyclostomes and elasmobranchs. Using histological and molecular techniques, we analyze the morphology and development of the joints of the little skate (*Leucoraja erinacea*), bamboo shark (*Chiloscyllium plagiosum*), and sea lamprey (*Petromyozon marinus*) as representatives of elasmobranchs and cyclostomes to find the origin of synovial joints in the phylogenetic tree of extant vertebrates. Early vertebrate evolution comprised multiple extinct lineages along the gnathostome stem [26]. Without soft-tissue preservation, whether an extinct group maintained synovial-like tissues is difficult to ascertain, but we review the literature to track fossil taxa that exhibited reciprocally articulated and cavitated bony joints and use this morphology as a proxy for the evidence of synovial joints. To identify the presence of the earliest synovial-like morphology in the evolutionary tree of vertebrates, we rule out anaspids, thelodonts, and galeaspids because of a lack of any reciprocal articulations in their skeletons [27, 28], and review the skeleton of jawless osteostracans and antiarch placoderms.

## Results and Discussion

### Cavitated joint articulations are present in chondrichthyans but absent in sea lampreys

To resolve the origin of synovial joints between chondrichthyans and cyclostomes, we analyze the skeletal morphology of late-stage embryos of the little skate (stage 33, *Leucoraja erinacea*) and juvenile sea lamprey (*Petromyozon marinus*). Stain enhanced micro-CT scanning reveals the presence of reciprocal articulating elements separated by a cavity in the skeleton of the little skate (Fig 1A-D), but the discrete cartilaginous elements in the sea lamprey are non-reciprocal and non-cavitated (Fig 1E-G), as evidenced through the presence of contrast in virtual sagittal sections. Histological analyses reveal the joint cavities in the jaw and pelvis of juvenile skates (Fig 1H,I), but no such cavity exists between the cartilaginous elements of the chondrocranium in the sea lamprey (Fig 1J,K).

**Fig 1.**
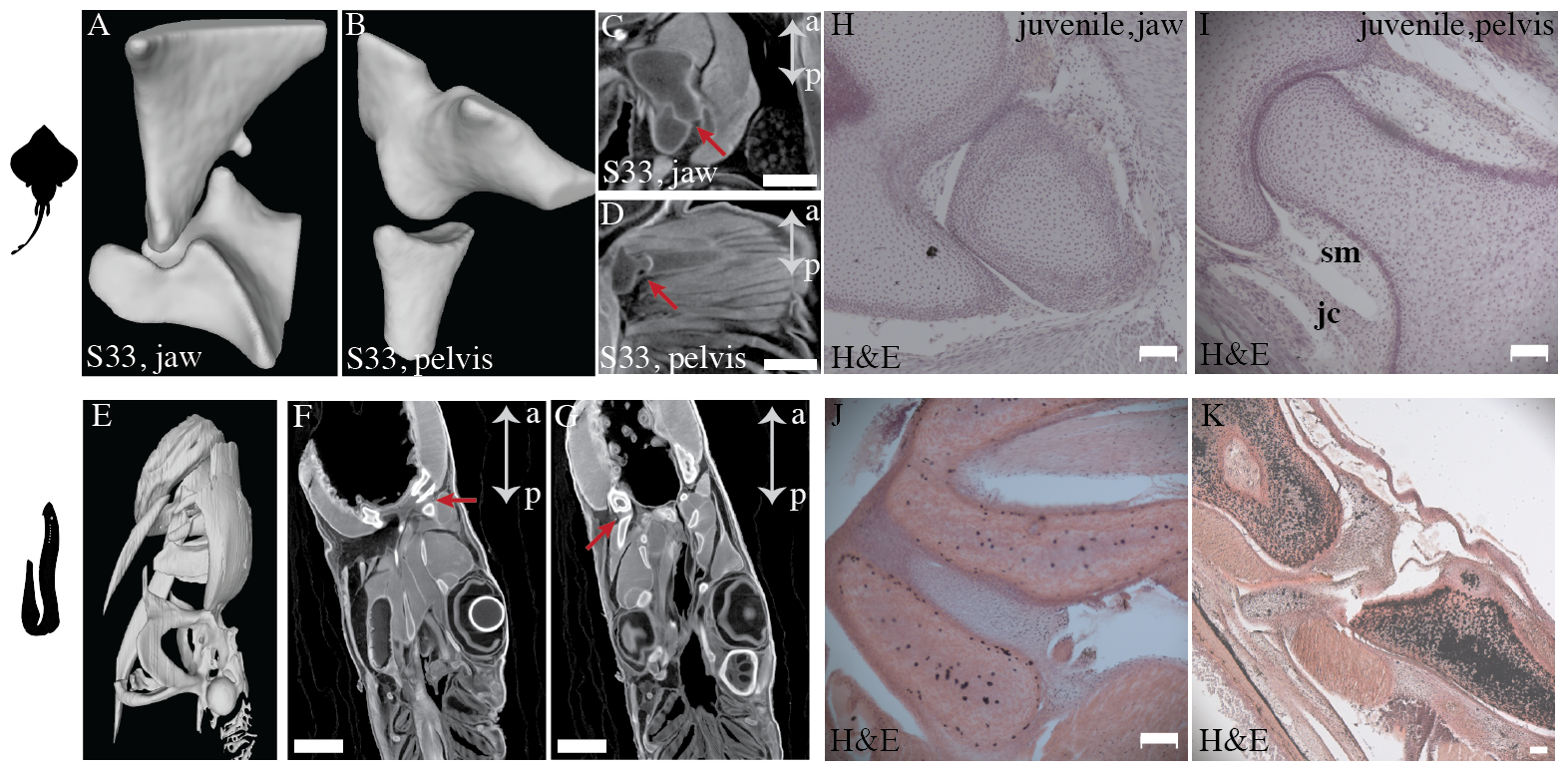
Micro-CT scans and histology show cavitated and reciprocal articulations in little skates (*Leucoraja erinacea*) but not in sea lampreys (*Petromyozon marinus*). (A, B) 3D reconstruction of the cartilaginous skeleton from the micro-CT scan of the little skate (stage 33) shows articulated joints with reciprocal surface geometries in the jaw (A) and the pelvis (B). (C, D) Image slices obtained from the micro-CT scans of the little skate display the articulations in the jaw (C), and the pelvic fin between the pelvic girdle and the first ray (D). (E) 3D reconstruction of the cartilaginous skeleton from the micro-CT scan of a juvenile lamprey in a lateral view. (F, G) Image slices obtained from the micro-CT scan of the juvenile sea lamprey display non-reciprocal cartilaginous elements in the chondrocranium and an absence of joint cavities, as evident by the presence of contrast between the neighbouring elements. (H, I) Histochemical staining show the presence of cavitated joints in the jaw (H) and the pelvis (I) of the juvenile skate. (J, K) Histochemical staining show the presence of uncavitated joints in the chondrocranium of the juvenile lamprey. Scale bars on the micro-CT scanned slices, 1) little skate: 0.5 mm, 2) lamprey: 1 mm. Scale bars on histology images: 100 µm. sm: synovial membrane, jc: joint capsule, a: anterior, p: posterior.

The joint cavity in the pelvis of the little skate at the juvenile stage is lined by a membrane characterized by a relatively acellular connective tissue containing folds, and an outer layer characterized by fibrous tissue connecting the two cartilaginous elements (Fig 2A). The structure of the lining resembles the synovial membrane and joint capsule found in tetrapods [29]. In little skates, a high density of flattened articular chondrocytes line the articular cavity and hypertrophic chondrocytes are present in the subarticular regions (Fig 2B). In sea lamprey, hypertrophic articular chondrocytes are evenly spaced through the cartilage elements and do not exhibit flattened morphology or change in density at the joint surfaces (Fig 1J,K). In a fixed whole mount skeleton of the little skate (stage 33) stained with Alcian Blue and Alizarin Red, manual opening and closing of the jaw shows relative sliding of the articular surfaces, similar to how synovial joints operate (Fig 2E,F). The morphology of skate joints is similar to the cavitated joints of a chicken embryo (Fig 2G,H,H’), that are known to exhibit relative sliding. Analysis of the developmental stages reveals that the embryonic skate belonging to stage 32 has uncavitated joints (Fig 2I,J) but stage 33 has cavitated joints (Fig 2K,L) in the jaw and the pelvis, showing that cavitation occurs between stages 32 and 33.

**Fig 2.**
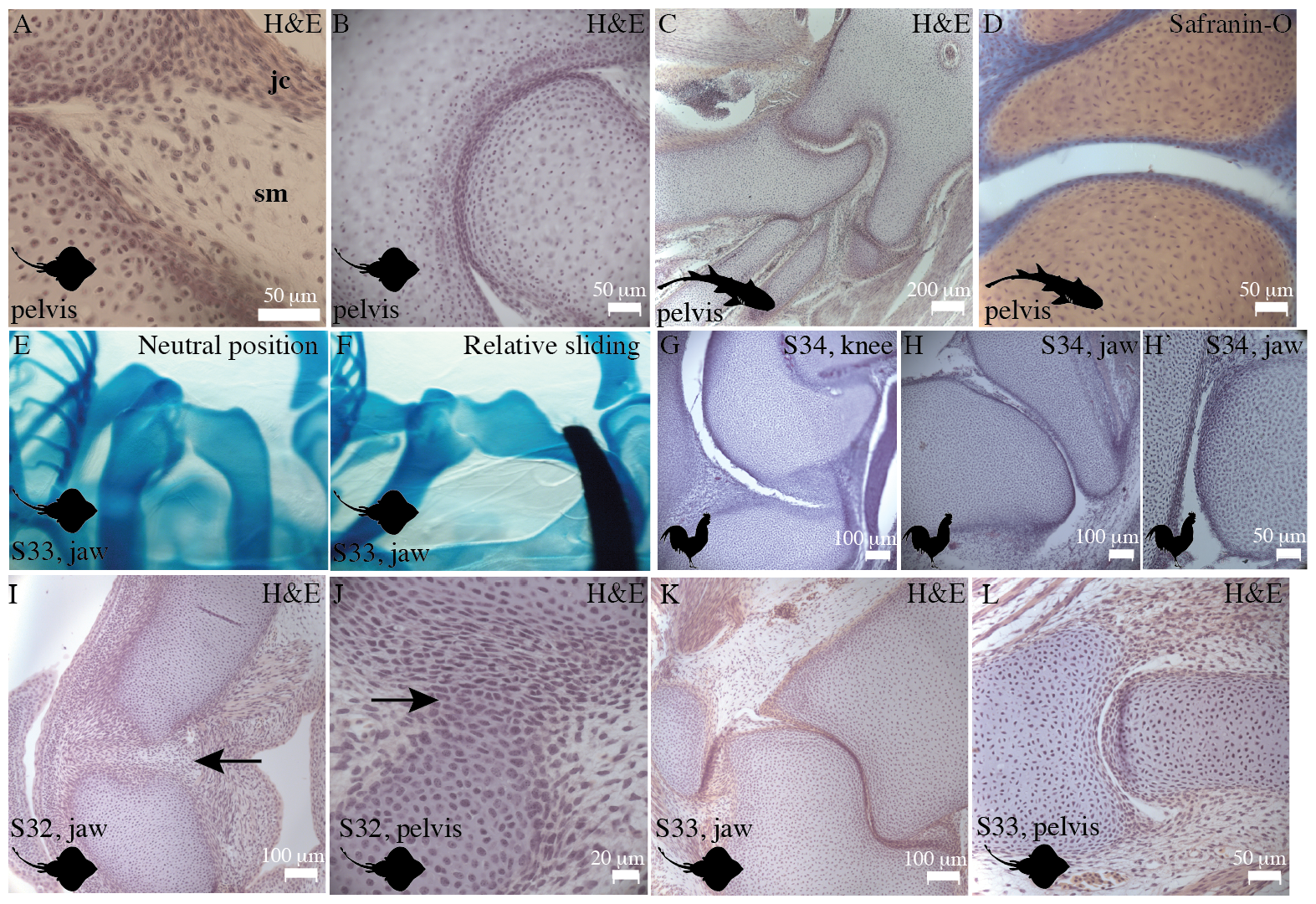
Comparison of the little skate and chicken joints exhibits similarities between chondrichthyan and tetrapod joints. (A,B) Hematoxylin and Eosin (H&E) histological staining of a little skate pelvis (juvenile) shows that individual cartilage elements in the pelvic fin are separated by a cavity. A distinct synovial membrane and joint capsule are present, and the joint capsule is in continuity with the perichondrium (A), and the chondrocytes at the articulations are flattened (B). (C,D) H&E and safranin-O staining of the sections of a bamboo shark, belonging to the group selachians, demonstrates the presence of cavitated joints and articulations lined by flattened chondrocytes in the pelvis. (E,F) Manual movement of the jaw in a whole mount Alcian and Alizarin Red stained specimen of the skate embryonic stage 33 demonstrates relative sliding of the articular surfaces (F) compared to the neutral position (E). (G,H,H’) H&E staining shows that the cavitated joints in the knee and the jaw of a chicken stage 34 embryo exhibits cavitated joints and flattened chondrocytes. (I,J) Sections of the jaw (I) and the pelvic joint (J) of an embryonic little skate (stage 32) stained with H&E display uncavitated joints and the presence of interzone (arrows). (K,L) Sections of the jaw (K) and the pelvic joint (L) of an embryonic skate (stage 33) stained with H&E display cavitated joints. sm: synovial membrance, jc: joint capsule.

To find whether cavitated joints are a conserved feature of elasmobranchs, we analyzed the joints of bamboo shark, belonging to selachians that is a sister group of batoids, and show the presence of cavitated joints with flattened chondrocytes lining the articular surfaces, showing a synovial-like morphology (Fig 2C,D). The presence of reciprocally cavitated articular surfaces containing flattened articular chondrocytes, lined by a synovium are evidence of synovial-like morphology in the little skate and bamboo sharks. Relative sliding of the articular surfaces, and the cavitation of uninterrupted mesenchyme to give rise to articulations in the little skate further show features of synovial-like development. Thus, morphological analyses suggests that cavitated synovial-like joints are a conserved feature in extant jawed vertebrates.

### Collagen-II forms the primary component of the subarticular cartilage and Collagen-I is limited to the articular surface in the chondrichthyan joints

Tetrapod articular cartilage is rich in collagen-I and collagen-II (Fig 3A, [5]), and although previous studies show that collagen-II forms the bulk of the cartilaginous skeleton of skates [19], the composition of the articular cartilage and its similarity to tetrapod hyaline cartilage remains unknown. We analyze the organization of the extracellular matrix in the articular and the subarticular cartilage in chondrichthyans by using the techniques of immunostaining for collagen-II and picrosirius staining for collagen-I. Our immunostaining data reveal that the articular surfaces as well as the subarticular cartilage in the cavitated pelvic joints of the skate embryo (stage 33), consist of collagen-II (Fig 3B,C). In the uncavitated joints belonging to stage 32 of the little skate, we find that the interzone consists of collagen-II; however, it is not as dense as the collagen-II present in the surrounding extracellular matrix and shows signatures of being mechanically strained (Fig 3D,E), likely from cyclic muscle contraction [12, 13]. Analysis of picrosirius red staining with a polarizer reveals that the articular surfaces and the joint capsule of the cavitated joints such as the jaw and the pelvis (Fig 3F,G) in the skate embryos (stage 33) consist of collagen-I, similar to tetrapods, such as chickens (Fig 3H).

**Fig 3.**
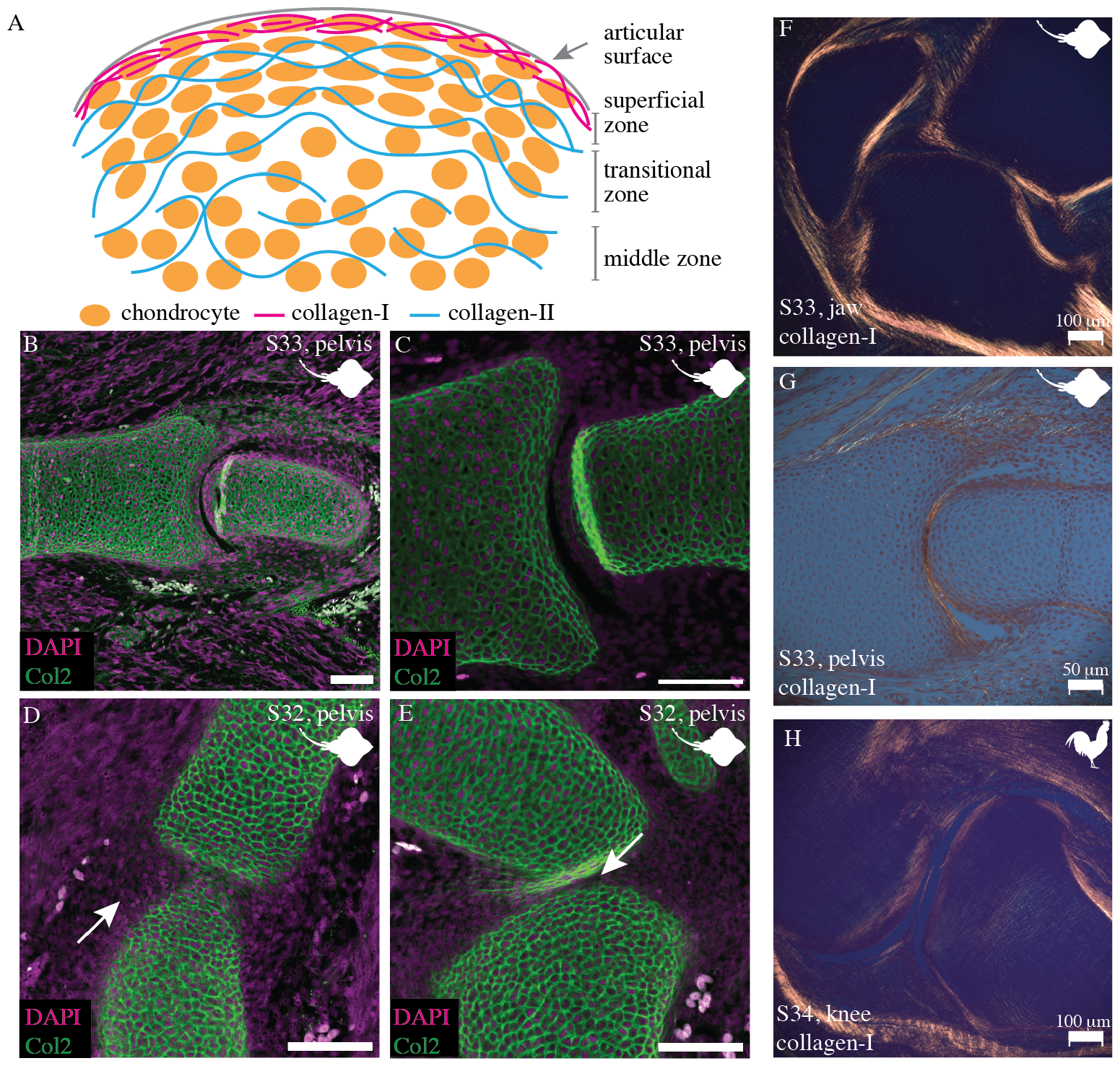
The subarticular surface of the little skate expresses collagen-II and the articular surface expresses collagen-I. (A) A pictorial representation of the organization of chondrocytes and ECM at the articular and the subarticular surfaces of a synovial joint. The superficial zone displays collagen-I, the transition zone displays collagen-II parallel to the articular surface, and the middle zone displays a random arrangement of collagen-II fibers. (B,C) The cavitated pelvic fin joints of the skate (stage 33) express collagen-II parallel to the articular surface and randomly distributed at the subarticular surface. (D,E) The uncavitated pelvic fin joints of the little skate (stage 32) display an interzone containing collagen-II that shows signs of being mechanically strained. (F-G) Collagen-I visualized in orange and yellow using picrosirius red staining is present at the articular surfaces and the perichondrium of the jaw (F) and the pelvic joint (G) of the little skate (stage 33). (H) The expression of collagen-I in the knee joint of a chicken embryo (stage 34) shows its expression at the articular surfaces and the perichondrium. Scale bars for immunofluorescence panels: 100µm.

### Sea lampreys and chondrichthyans display glycosaminoglycans and proteoglycans at the articular surfaces in a pattern similar to tetrapods

Glycosaminoglycans, polyanionic extracellular matrix molecules like hyaluronic acid, chondroitin sulfate, and keratan sulfate [30] present in the articular cartilage of tetrapods, attract water and keep the tissue hydrated to provide lubrication and shock absorption capacity to the synovial joints. Glycosaminoglycans such as hyaluronan [31] covalently attach to core proteins such as aggrecan [6] and lubricin [2] to form proteoglycans, critical for lubrication and load bearing function of the articular cartilage (Fig 4A, [32]). Earlier studies show that proteoglycan is expressed in the vertebral cartilage of chondrichthyans [33, 34], in the notochord sheath of lampreys [35], and in the cartilage of lamprey embryos and larvae [36]. Using histochemical staining and immunostaining, we analyze the presence and distribution of GAGs and proteoglycans in the cartilaginous skeleton at the joints in the little skates and juvenile lampreys.

**Fig 4.**
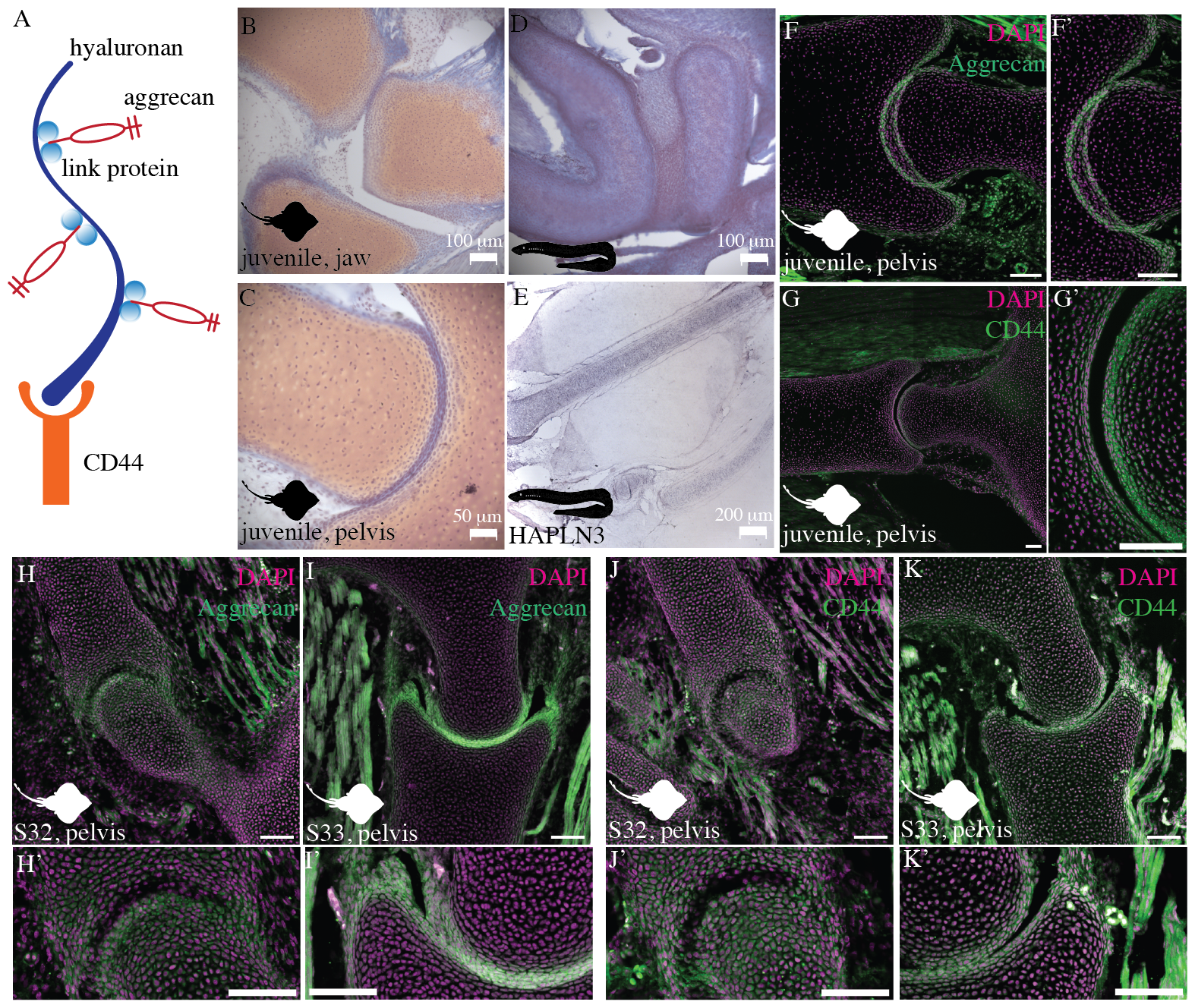
Little skate and the sea lamprey cartilage display glycosaminoglycans (GAGs), proteoglycans, and glycoproteins. (A) A pictorial representation of the interaction between GAGs such as hyaluronan, link proteins such as HAPLN3, and proteoglycans such as aggrecan. (B-D) Safranin-O staining shows the presence of glycosaminoglycans in the jaw (B) and pelvis (C) of the little skate, and in the pharyngeal cartilage of the sea lamprey (D). (E) *in situ* hybridization reveals that the cartilaginous skeleton of the sea lamprey expresses the proteoglycan linking protein, HAPLN3. (F,F’) The pelvic joint of a juvenile skate expresses aggrecan, a proteoglycan specific to synovial joints, at the articular surfaces. (G,G’) CD44, a cell surface glycoprotein and hyaluronic acid receptor, is expressed at the articular surfaces and the subarticular regions of the pelvic joints in the juvenile skate. (H-I’) The little skate expresses aggrecan at the interzone and the subarticular cartilage in the uncavitated stage 32 (H,H’) and cavitated stage 33 (I,I’) embryos. (J-K’) The skate expresses CD44 at the interzone and in the subarticular cartilage in the uncavitated stage 32 (J,J’) and cavitated stage 33 embryos (K,K’). Scale bars for immunofluorescence panels: 100µm.

Using safranin-O staining, we find that the cartilage of juvenile little skates is rich in glycosaminoglycans (GAGs) (Fig 4B,C). Compared to previous studies that measured the amount of GAGs in the skate cartilage by using biochemical assays [37], we focus on the distribution of GAGs near the articular cartilage. In the jaw and pelvic joints of the skate, the amount of GAGs is lower at the articular surface compared to the deeper cartilage (Fig 4B,C), similar to tetrapods. Sea lamprey cartilage has GAGs throughout the skeleton, mostly concentrated in the middle of the cartilage elements (Fig 4D).

Using mRNA *in situ* hybridization, we find that juvenile sea lamprey expresses hyaluronan and proteoglycan link protein (HAPLN3) throughout the cartilage skeleton such that the expression is not limited to the joint surfaces (Fig 4E). Hyaluronan and proteoglycan link protein HAPLN3, known to help form proteoglycan aggrecates [6, 38, 39] (Fig 4A), organizes and stabilizes the interaction between hyaluronic acid and the extracellular matrix [40]. In comparison, immunostaining of juvenile skate with cavitated joints reveals that the expression of proteoglycan aggrecan is limited to the articular surfaces (Fig 4F,F’,Fig S1). Aggrecan synthesis at the articular surfaces is mechanosensitive, and aggrecan interacts with hyaluronic acid to give rise to a proteoglycan structure that improves the shear and compressive load-bearing capacity of the cartilage [6,41,42]. Immunostaining for CD44, the principal cell surface glycoprotein that acts as a receptor for hyaluronic acid [43], shows that CD44 is expressed at the articular surfaces as well as in the subarticular regions, albeit at lower concentrations, in the juvenile skate (Fig 4G,G’, Fig S1). In addition, both aggrecan and CD44 are also expressed in the synovial membrane, consistent with their expression in tetrapods [29]. The expression of proteoglycans and glycoproteins in juvenile chondrichthyans is consistent with their role in articular surface lubrication.

The embryonic stage 32 of little skates with uncavitated joints expresses aggrecan throughout the cartilaginous skeleton (Fig 4H,H’,Fig S1), as shown previously [19], whereas stage 33 shows a greater expression at the articular surfaces compared to the bulk of the cartilage (Fig 4I,I’,Fig S1). Hyaluronan and CD44 play a role in synovial cavity formation during development [43–46]. Immunostaining for CD44 reveals its expression in the interzone as well as the bulk cartilage of the joints of the stage 32 little skate (Fig 4J,J’), and a greater expression at the articular surfaces at the cavitated joints of stage 33 little skate (Fig 4K,K’). Although aggrecan becomes localized to the articular surfaces in juvenile skates (Fig 4F,F’), CD44 is present in the bulk of the cartilage (Fig 4G,G’). Whether adult skates exhibit a localized expression of CD44 at the articulations similar to tetrapods [44] remains unknown. The expression of proteoglycans and glycoproteins in the joints of embyronic chondrichthyans is consistent with their role in articular surface development.

### Molecular mechanisms underlying chondrichthyan joint development are similar to tetrapods

Next, we examine whether joint morphogenesis in the little skate relies on molecular mechanisms similar to those in tetrapods. To this end, we analyze the expression of key proteins from two signaling pathways, growth differentiation factor 5 (*Gdf5*) from the BMP signaling pathway [7, 47] and *β*-catenin from the Wnt signaling pathway [8]. Using *in situ* hybridization, we find that the skate at embryonic stage 32 expresses *Gdf5* at the presumptive interzones during development (Fig 5A-D). However, the expression is not limited to cavitated joints (Fig 5A,B,C) but is also present in the non-cavitating intervertebral joints (Fig 5D). Immunostaining reveals that the pelvic joint of the little skate belonging to stage 32 and stage 33 express *β*-catenin at the the articular surfaces (Fig 5E,E’,G) but the expression is not noticeable in the intervertebral discs (Fig 5F).

**Fig 5.**
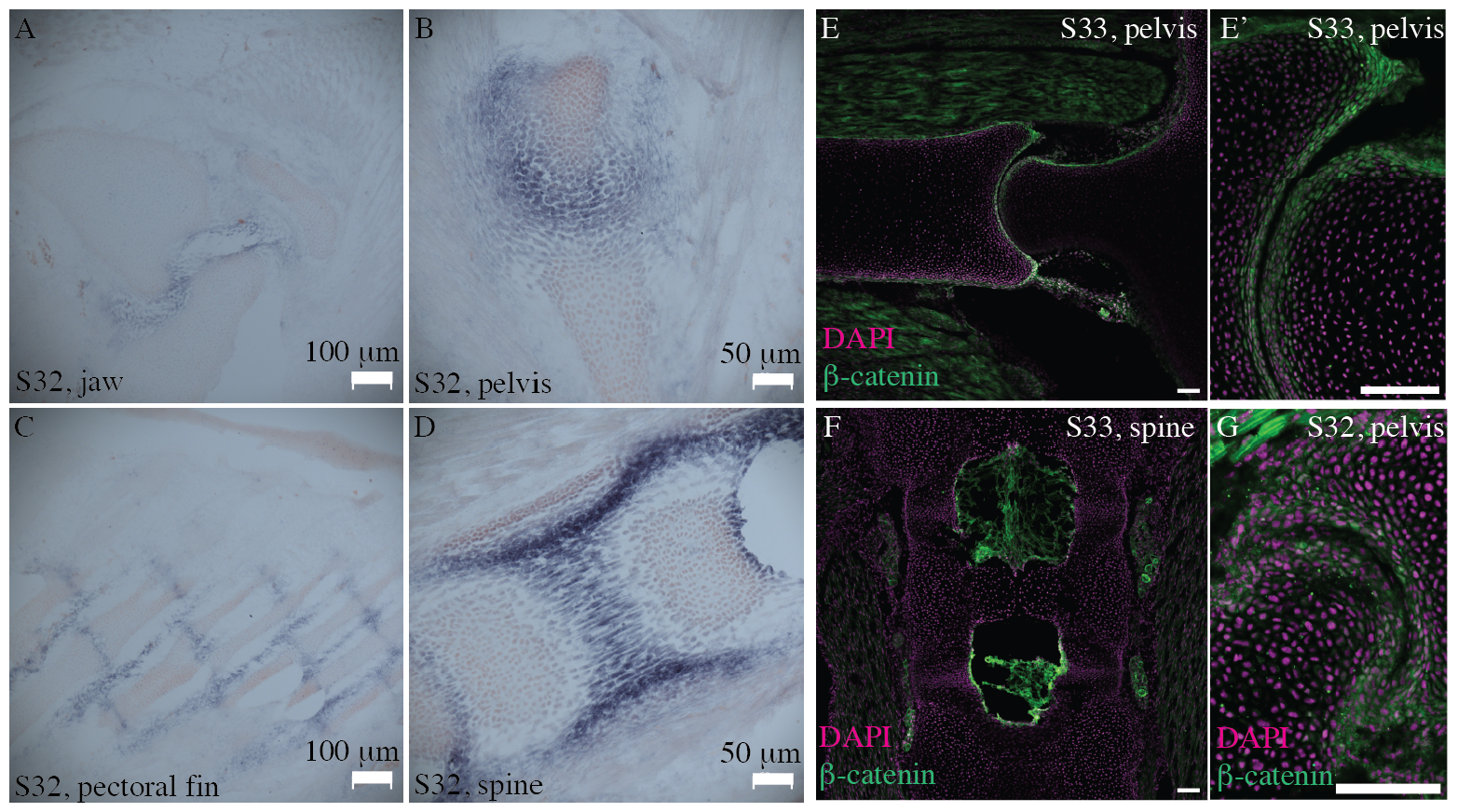
BMP and Wnt signaling play a role in the joint morphogenesis in the little skate. (A-D) *in situ* hybridization reveals that growth differentiation factor 5 (*Gdf5*) is expressed at the joint interzones in the embryonic little skate at stage 32. The expression is seen at the interzones of the jaw (A), pelvis (B), and pectoral fin (C) joints that develop to form a joint cavity, and also in the intervertebral discs (D) that do not cavitate. (E-F) Immunostaining shows the expression of *β*-catenin in stage 33 embryos of the little skate at the articular surfaces of the pelvic joint (E,E’), but not in the intervertebral joints (F). (G) The uncavitated pelvic joint of the little skate (stage 32) expresses *β*-catenin at the articular surfaces. Scale bars for immunofluorescence panels: 100µm.

### Muscle contraction is essential for joint cavitation in the embryos of the little skate

By inducing paralysis in the embryonic little skates from stage 31 for twelve days and stage 32 for seven days by using the anesthetic tricaine mesylate (MS-222) and comparing them with identically aged non-paralyzed embryos, we tested the effect of muscle contraction on joint cavitation. The comparison of control and paralyzed stage 31 little skate embryos shows that control embryos are further along the process of cavitation, demonstrated by flattened articular chondrocytes in the interzone and a better definition of articular geometry (Fig 6A-C) compared to the paralyzed embryos (Fig 6A’-C’). Histological comparison of paralyzed with the control stage 32 embryos reveals that control animals exhibit cavitated joints (Fig 6D-F), and the identically aged but paralyzed embryos lack a clear joint cavity (Fig 6D’-F’). Because Wnt14-signaling is mechanosensitive [13], we compare the expression of *β*-catenin in paralyzed versus control embryos to find that control embryos express higher amounts of *β*-catenin in the chondrocytes at the articular surface compared to the paralyzed embryos for both stage 31 (Fig 6G-H’) and stage 32 (Fig 6I-J’). Wnt-signaling plays a role in the downstream regulation of the expression of CD44 (hyaluronic acid receptor) at the articular surfaces [9], and the mechanosensitive secretion of hyaluronic acid [48] plays a role in the process of joint cavitation [45, 46]. Thus, our investigations of development of joints in the little skate suggest that their jaw and pelvic joints exhibit a synovial morphology.

**Fig 6.**
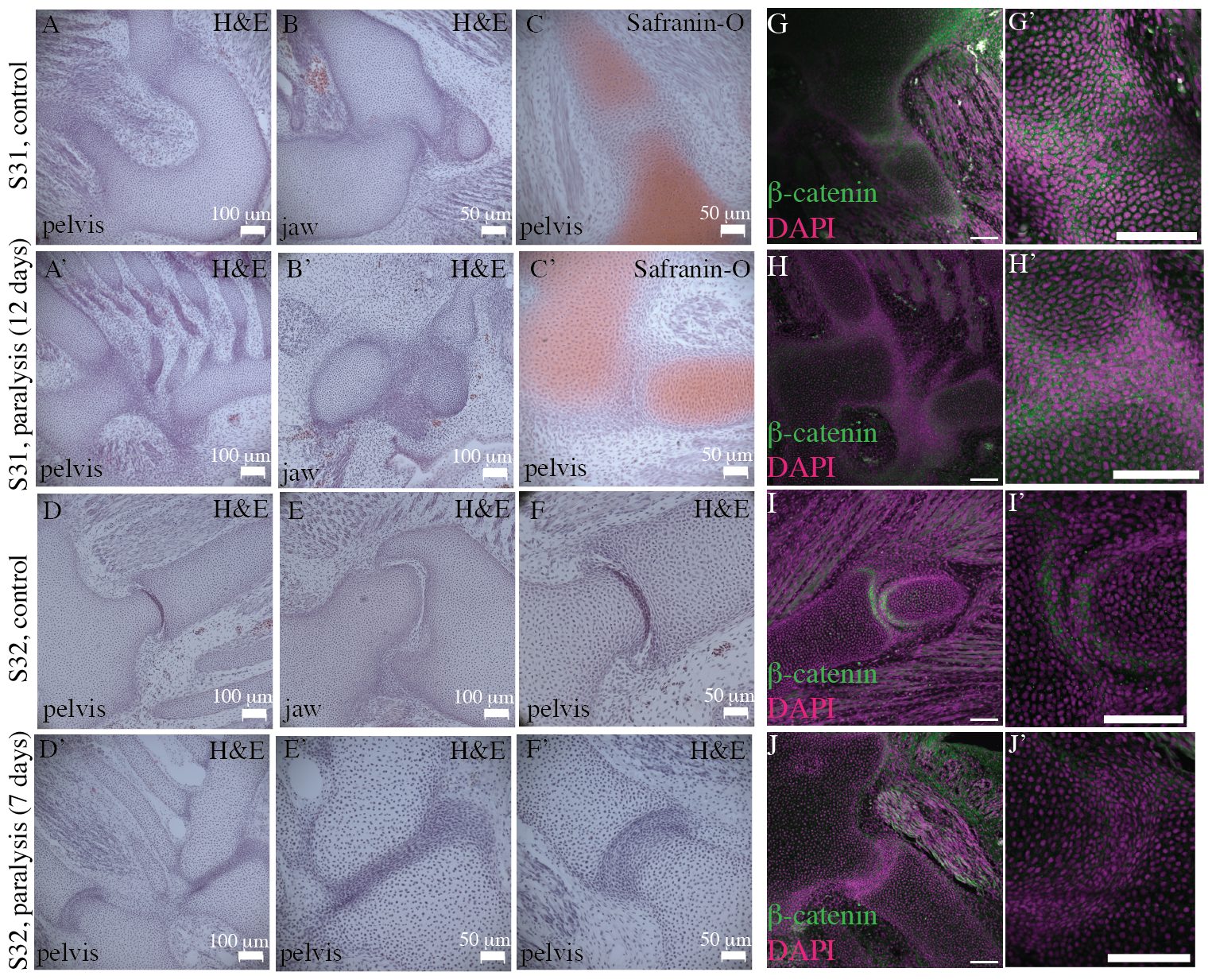
Muscle paralysis leads to joint fusion and impairs *β*-catenin signaling in the little skate. (A-C) Histological staining using H&E and safranin-O of the little skate (stage 31) control embryos raised for 12 days in sea water show that the joints are not fully cavitated in the pelvis (A,C) and the jaw (B) but the interzone shows the definition of the developing articular surfaces. (A’-C’) Histological staining using H&E and safranin-O of the little skate (stage 31) embryos raised in sea water infused with tricaine mesylate to induce paralysis shows a broader interzone and undefined articular surfaces compared to the control embryos. (D-F) Histological staining using H&E of stage 32 skate control embryos raised for 7 days in sea water shows the presence of distinct joint cavities in the jaw and the pelvic fin. (D’-F’) Histological staining using H&E of stage 32 skate embryos raised for 7 days in sea water infused with tricaine mesylate to induce paralysis shows that pelvic joints are fused and uncavitated. (G-H’) Immunostaining for *β*-catenin in the stage 31 little skate embryos shows that it is expressed in the articular cartilage of both the control and the paralyzed embryos, but a comparatively uncharacteristic and lower expression in the paralyzed embryos. (I-J’) Immunostaining for *β*-catenin in the stage 32 little skate embryos shows expression at the articular surfaces of the control embryos, but not in the paralyzed embryos. Scale bars for immunofluorescence panels: 100µm.

During the paralysis experiments, we used tricaine mesylate and let the embryos develop inside their egg cases, leading to difficulties in the interpretation of our experimental results. Although egg cases are not entirely closed and allow movement of surrounding aerated water, skates are additionally known to rely on their tail movement to allow the aerated water inside the egg cases. We attempted to remedy this by additional oxygenation, but it cannot be ruled out that paralysis led to lower oxygen inside the egg cases, leading to stunted growth (Fig S2). Tricaine mesylate likely works by inhibiting neural voltage-gated sodium channels at low concentrations, and inhibits evoked muscle contraction at high concentrations by a yet unknown mechanism, and its pharmacological side effects on development are not clear [49]. Thus, the absence of cavitation may not be a direct consequence of muscle paralysis, and fused joints could be a consequence of the pharmacological side effects due to general anesthesia. However, our results are consistent with experiments performed on tetrapods such as chickens and mice, that also exhibit stunted growth and fused joints [11, 16, 50].

## Conclusion

Our data demonstrate that elasmobranch joints consist of reciprocally articulating surfaces separated by a cavity. Expression of *Gdf5, β* -catenin, proteoglycans, and glycoproteins at the cavitated joints of chondrichthyans suggest that cartilaginous fishes and bony fishes rely on similar molecular regulatory mechanisms and proteins for the morphogenesis and function of synovial joints. Thus, our histological and molecular investigations show considerable morphological and developmental similarities between chondrichthyan and osteichthyan joints (Fig 7A). We conclude that chondrichthyans consist of synovial joints, but cyclostomes like sea lamprey do not. The discovery of synovial joints in chondrichthyans challenges the view that the phenotype of synovial joints is associated only with bony fishes. The results here indicate that synovial joints evolved before the divergence of cartilaginous and bony fishes. In conclusion, synovial joints are a shared feature of extant jawed vertebrates (Fig 7B).

**Fig 7.**
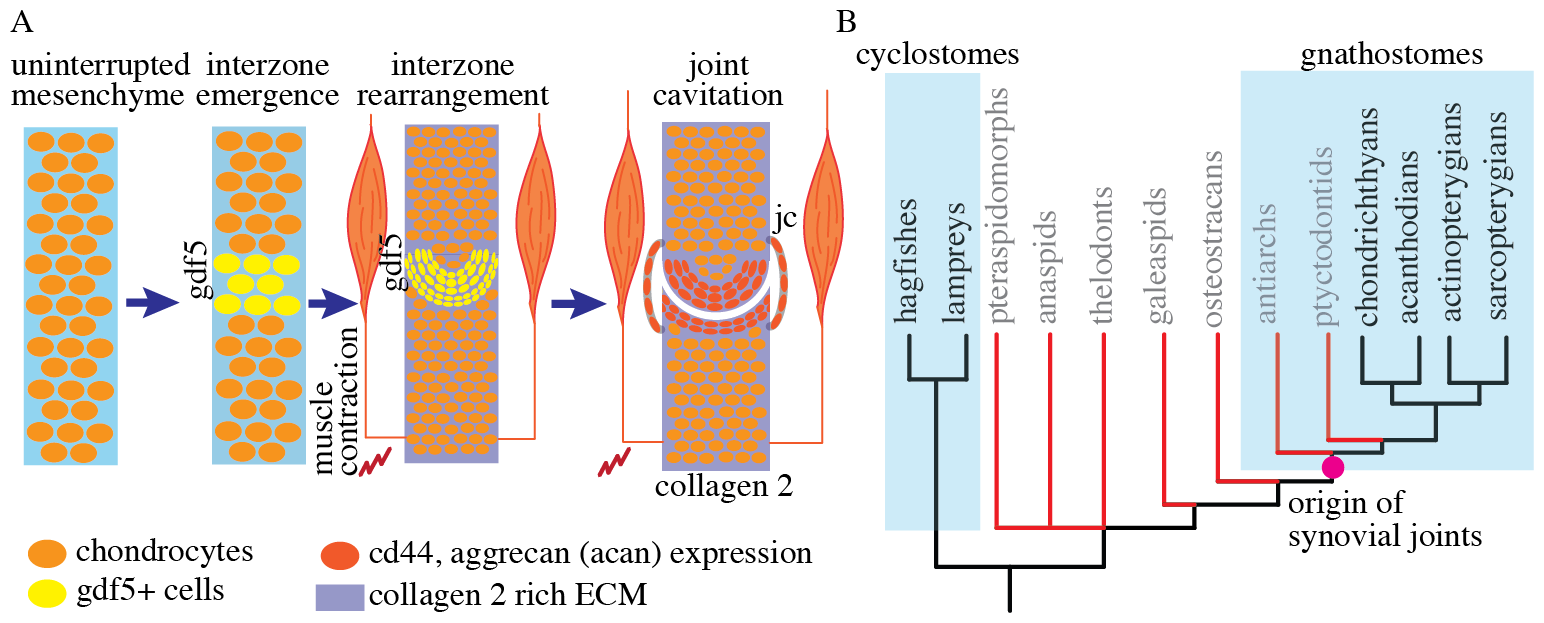
Synovial joints are present in elasmobranchs and evolved in the common ancestor of gnathostomes. (B) A pictorial representation of the key mechanisms required for the morphogenesis of synovial joints in chondrichthyans. Joint development relies on the emergence of interzone in the uninterrupted mesenchyme that cavitates to form a joint cavity by relying on BMP and Wnt signaling, and muscle contraction. jc: joint capsule. (B) Our results provide evidence that synovial joints exist in extant jawed vertebrates (gnathostomes) but not in cyclostomes, and the presence of cavitated joints in antiarchs suggests that joints that function by sliding like synovial joints originated in the common ancestor of gnathostomes. Phylogenetic tree adapted from Donoghue and Keating, 2014 [26].

Our results show that the regulatory mechanisms required for the morphogenesis of synovial joints such as *Gdf5* are also expressed in non-cavitating intervertebral joints of the skate. Studies show that *Gdf5* is expressed in sea lamprey in the pharyngeal arches and mucocartilage of the ventral pharynx [21]. If future studies reveal that lampreys also express *Gdf5* at their prospective cartilaginous joints, it would suggest a conserved role of BMP pathway in the development of cartilaginous and synovial joints in vertebrates. Furthermore, proteoglycans and GAGs are not limited to articular cartilage, but are expressed throughout the cartilaginous skeleton of sea lampreys as suggested by the expression of hyaluronan binding proteins and glycosaminoglycans. The expression of proteoglycans and GAGs in the lamprey cartilage suggests their primitive role in regulating osmotic and turgor pressure in the cartilaginous skeleton, likely for countering the mechanical trauma in the absence of mineralization [39], similar to its function in chondrichthyans [37, 51]. Therefore, we hypothesize a co-option model wherein the molecular pathways required for the development and function of synovial joints were derived from the novel recruitment of pre-existing BMP and Wnt signaling pathways, and the expression of proteoglycans at the articulations.

Chondrichthyans were earlier hypothesized not to have synovial joints because their flexible cartilaginous skeleton obviates the necessity for a joint structure that improves the range of motion [3]. However, the mineralized as well as non-mineralized cartilaginous skeleton of elasmobranchs can be as stiff and strong as mammalian trabecular bone owing to its collagen and glycosaminoglycan content [37, 51, 52], suggesting that their bendability is not superior to bony skeletons. In comparison, lamprey fibrocartilage compares with mammalian articular cartilage in its mechanical properties [53], and therefore, it is comparatively softer than chondrichthyan cartilage. Lamprey cartilage also expresses lamprin, an elastin-like protein, in their nasal, branchial, and pericardial cartilage, imparting enhanced sketelal flexibility [54]. Our results are consistent with the functional studies in showing that chondrichthyans, like tetrapods, rely on synovial joints for exhibiting considerable jaw and fin movements in their stiff skeleton that reaches several meters in size, but to the exception of rest of the vertebrates, lampreys do not.

Finally, we review the literature on cavitated joints in the extinct lineages along the gnathostome stem, focusing on the jawless osteostracans and stem gnathostome placoderms. In osteostracans, debates exist on whether the endoskeletal pectoral fins made up of calcified cartilage articulated with the exoskeleton of the shoulder girdle [27, 55–57]. We note that the articular surface of the shoulder girdle in osteostracans such as *Cephalaspis* consisted of multiple canals and fenestrae to allow passage for muscles and nerves (see Fig 5, 16, 17, 18 in [57]), suggesting that the articulation was cavitated; however, the evidence is not definitive. In antiarch placoderms such as *Bothriolepis canadensis*, morphological analyses suggest the presence of cavitated joints in the pectoral fins [58, 59]. Furthermore, the hypothesized functional ranges of motion in the fin of *Bothriolepis canadensis* are consistent with relative sliding of reciprocally shaped articular surfaces [58], similar to how synovial joints generate motion in tetrapods. Therefore, we conservatively estimate the minimal occurrence of cavitated joints at the common ancestor of gnathostomes (Fig 7B). Compared to osteostracans that were comparatively smaller [60], placoderms at their extremes consisted of predatory *Dunkleosteus* reaching greater than 5 meters in length [61] and exhibited a great diversity in ecological niche occupation [62]. Whether cavitated joints contributed to their greater size and predatory performance compared to coexisting fishes with only cartilaginous or fibrous joints, due to improved feeding and locomoting abilities is a question for future investigations.

## Materials and methods

### Animal procurement, care, and procedures

The Institutional Animal Care and Use Committees of the University of Chicago approved the procedures on the little skate and sea lamprey (IACUC# 71033-05). All the procedures were performed at the University of Chicago. The juvenile and embryonic stages 31 (16-26 weeks old), 32 (27-29 weeks old), and 33 (*>* 29 weeks old) [63], and juvenile bamboo sharks were obtained from the Marine Resources Center, Marine Biological Laboratory, Woods Hole, MA, USA. Juvenile sea lampreys were obtained from Acme Lamprey (Harrison, ME, USA). For micro-CT scanning, histology, immunostaining, *in situ* hybridization, and after the completion of the muscle paralysis experiments, the animals were euthanized using 0.5% tricaine methanesulfonate (MS-222, Syndel) until the cessation of heartbeat and fixed in 4% PFA (Acros Organics, Item No. EW-88353-82) overnight (skate embryos), or for four days (juvenile sea lamprey and skates) before transferring them to 1X PBS (Fisher Bioreagents).

### Micro-CT scanning

Scanning was performed on Phoenix v|tome|x S 240 from GE (PaleoCT facility, RRID: SCR024763, University of Chicago) using the 180 kV nano-focus tube. Sea lamprey was stained using Ruthenium red (Cayman Chemical, CAS Number 11103-72-3) to give rise to contrast between cartilage and muscles following the protocol of Gabner et. al [64] with the modification that the juvenile lampreys were kept in the staining solution for seven days. The scanning parameters were 80 kV of Voltage, 180 µA beam current, and images were generated with a voxel size of 14.238 µm. The skate was stained in phosphomolybdic acid (Sigma Aldrich). Specimens were stepped in 25%, 50%, and 75% solutions of sucrose (Sigma Aldrich) in 1X PBS for one hour each with shaking, and then transferred to 20% sucrose in PBS overnight. The embryos were transferred to 5% PMA solution in 1X PBS for a week before scanning. The scanning parameters used were 90 kV Voltage, 270 µA beam current, and the images were generated with a voxel size of 8.738 µm. Segmentation and reconstruction were performed on Amira 3D 2021.1 (©1995-2001 Konrad-Zuse-Zentrum Berlin (ZIB), ©1999-2021 FEI SAS, a part of Thermo Fisher Scientific). MeshLab v2023.12 [65] was used for visualization.

### Histochemical staining

The embryonic and juvenile little skates (*Leucoraja erinacea*), sea lamprey (*Petromyozon marinus*), bamboo shark (*Chiloscyllium plagiosum*), and chicken (*Gallus gallus*) were stepped in 70% ethanol and paraffin embedded. Embedding was performed by the Human Tissue Resource Center at the University of Chicago. The paraffin-embedded blocks were sectioned at 10µm using a rotary microtome (HM 330, Microm Heidelberg). Sections were mounted on TOMO IHC adhesive glass slides (TOM-1190, Matsunami). H&E staining was performed using Hematoxylin (Sigma-Aldrich) and Eosin (Sigma-Aldrich) following the protocol of Gillis et. al, 2009 [66]. Safranin-O staining was performed using Safranine-O (Fisher-Scientific), Hematoxylin, and Fast Green FCF (Sigma Life Science). The sections were dewaxed and hydrated to distilled water, stained with Weigert’s iron hematoxylin working solution for 10 minutes, washed in running tap water for 10 minutes, stained with fast green (FCF) solution for 5 minutes, rinsed in 1% acetic acid solution for 15 seconds, stained with 0.1% safranin O solution for 5 minutes, and stepwise dehydrated in increasing concentrations of ethanol and finally, histosol. Picrosirius staining was performed using Direct Red 80 (Sigma Aldrich). Slides were dewaxed and hydrated to distilled water, stained with Weigert’s iron hematoxylin working solution for 10 minutes, washed in running tap water for 10 minutes, stained in picrosirius red solution (0.1% Direct Red 80 in saturated aqueous solution of picric acid) for an hour, washed in 0.5% acetic acid solution, and dehydrated in 100% ethanol and histosol. Slides sections were mounted using Permount (FisherChemicals, Fisher Scientific).

### Immunofluorescence

Fixed tissues were stepped in 1x PBS overnight and then in 20% sucrose solution prepared in 1x PBS before embedding them in Tissue-Plus OCT compound (Fisher Healthcare). Cryosections were obtained at a thickness of 10µm using Leica CM3050 S. Apart from Collagen-II immunostaining that was performed on paraffin sections, immunostaining for aggrecan, CD44, and *β*-catenin was performed on cryosections. Paraffin sections of thickness 10µm were prepared as described above. Collagen-II immunostaining on paraffin sections was performed using the protocol of Gillis et. al, 2012 [67]. Aggrecan, CD44, and *β*-catenin immunostaining was performed by washing the cryosections in 1x PBS for 5 minutes, blocking them in blocking buffer (5% goat serum in PBT prepared with 0.1% Tween-20, Sigma Aldrich) for an hour at room temperature followed by overnight primary antibody incubation at 4°C. Primary antibody was washed 3 times in 1X PBS for 5 minutes each, followed by overnight incubation in secondary antibody, washed three times in 1x PBS for 5 minutes each the next day followed by mounting. Primary and secondary antibodies were diluted in blocking buffer. The following primary antibodies were used: Collagen-II (II.II6B3, Developmental Studies Hybridoma Bank, University of Iowa; 1:20), *β*-Catenin (ProteinTech, 17565-1-AP, 1:300), Aggrecan (12/21/1-C-6, Developmental Studies Hybridoma Bank, University of Iowa; 1:20), Hyaluronic acid receptor (H4C4, Gene name: CD44, Developmental Studies Hybridoma Bank, University of Iowa; 1:20). The following secondary antibodies were used at a dilution of 1:300: Cy3 donkey-anti-rabbit (Jackson, 711-166-152), Cy5 donkey-anti-mouse (Jackson, 715-175-151), CF 633 goat-anti-mouse (Biotium, 20120-1), CF633 goat-anti-rabbit (Biotium, 20122-1), and Cy3 donkey-anti-mouse (Jackson). The slides were mounted using Fluoromont-G with DAPI (Invitrogen). All immunostaining was carried out in quadruplets and the controls were prepared by following the same protocol but in the absence of the primary antibody.

### mRNA *in situ* hybridization

Probes were designed for *Petromyzon marinus* hyaluronic acid and proteoglycan link protein 3 (HAPLN3), containing a hyaluronan (HA)-binding domain found in CD44, using the sequences from National Library of Medicine, National Center for Biotechnology Information (GenBank XM032975677). For designing the probe, we amplified the cDNA of the sea lamprey using CCGAATAGTGTGGTCAGGGT as the forward primer and TGCGGCTTCATTTCATACGG as the reverse primer. The fixed juvenile sea lamprey was stepped in 100% methanol and kept at -20°C before it was paraffin embedded and sliced by a microtome into a thickness of 10 µm sections. *in situ* hybridization was performed on the paraffin sections following the protocol by Sugahara et. al, 2015 [68]with the modifications that we used 10 µg/ml of proteinase-K (Thermo Scientific) for 10 minutes for the digestion of sections, Nuclear Fast Red (Newcomer Supply) for counterstaining after the color reaction, and Permount (FisherChemicals, Fisher Scientific) as a mounting medium.

For *in situ* hybridization in the little skate for analyzing the expression of growth differentiation factor 5 (gdf5), we selected the sequence available for *Amblyraja radiata* (GenBank XP032897637) and performed a nucleotide BLAST against the little skate transcriptomic contigs [69, 70] and found a high degree of similarity with contig 44189 (bit score = 383, identity = 215/217 with a query length of 817, similarity = 99, e-value = 2e-105). We designed probes for contig 44189 by amplifying the cDNA of the little skate using TGCACTCTCCGATTGTCCAA as the forward primer and GCAACCACAGGACTCTACCA as the reverse primer. The fixed skate embryos were stepped in 100% methanol and stored at -20°C before paraffin embedding. For *in situ* hybridization on the paraffin sections, we followed the protocol in [71] with modifications according to [19, 67].

### Muscle paralysis

Little skates from the embryonic stages 32 (n=2) and 31 (n=2) were raised in rectangular tanks and the same number were maintained in control tanks. Water temperature was maintained at 17°C and aeration was achieved using air bubblers. The experimental tank consisted of 1500 ml of water mixed with 0.25 grams of tricaine methanesulfonate and 45 grams of sea salt (Instant Ocean, Spectrum Brands Pet LLC, Blacksburg, VA). The control tank contained 1500 ml of water and 45 grams of sea salt but no tricaine methanesulfonate. Water in both the tanks was changed every two days, and mortality was checked. For stage 32, we collected embryos from the control and the experimental group after seven days of the experiment. For stage 31, we collected the embryos after 12 days and 18 days of the experiment. We performed histology on the paraffin sections with Hematoxylin and Eosin and Safranin-O, and immunostaining for *β*-catenin, as described above.

### Microscopy and Image Analysis

Microscopy for histochemical staining was performed on Axioskop 2 mounted with Zeiss Calibri 7 camera and a polarizer. The software used for image acquisition was Zen 3.8 (Carl Zeiss Microscopy GmbH). Whole mount specimens were imaged on Leica DFC425 C and the software used for image acquisition was AmScope x64 (4.11.20896.20220521). Immunostaining slides were imaged at Zeiss LSM 710 Axio Imager 2 (Release Version 8.1.0.484, Carl Zeiss Microscopy GmbH) and the software used was Zen 2012. ImageJ2 Version 2.14.0/1.54f and Adobe Illustrator 28.0 (©1987-2023 Adobe) were used for processing images and constructing the plates, respectively.

## Acknowledgments

Shubin lab members including Sam Norris and Matteo Fabbri for help with immunostaining and microscopy, Melvin Bonilla for help with *in situ* hybridization, and Shiri Kult, Emily Hillan, Washaakh Ahmad, and Dylan Jockel for discussions and support. April Neander for help with microCT scanning. Tom Stewart for discussions and comments on the draft of the manuscript. Brinson Family Foundation, University of Chicago Biological Sciences Foundation, and HFSP - RGP0010/2022 for funding.

## Notes

### Competing Interest Statement

The authors have declared no competing interest.

